# Biological Reinforcement Learning via Predictive Spacetime Encoding

**DOI:** 10.1101/2020.08.21.260844

**Authors:** Minsu Abel Yang, Jee Hang Lee, Sang Wan Lee

## Abstract

Recent advances in reinforcement learning (RL) have successfully addressed several challenges, such as performance, scalability, or sample efficiency associated with the use of this technology. Although RL algorithms bear relevance to psychology and neuroscience in a broader context, they lack biological plausibility. Motivated by recent neural findings demonstrating the capacity of the hippocampus and prefrontal cortex to gather space and time information from the environment, this study presents a novel RL model, called spacetime Q-Network (STQN), that exploits predictive spatiotemporal encoding to reliably learn highly uncertain environment. The proposed method consists of two primary components. The first component is the successor representation with theta phase precession implements hippocampal spacetime encoding, acting as a rollout prediction. The second component, called Q switch ensemble, implements prefrontal population coding for reliable reward prediction. We also implement a single learning rule to accommodate both hippocampal-prefrontal replay and synaptic homeostasis, which subserves confidence-based metacognitive learning. To demonstrate the capacity of our model, we design a task array simulating various levels of environmental uncertainty and complexity. Results show that our model significantly outperforms a few state-of-the-art RL models. In the subsequent ablation study, we showed unique contributions of each component to resolving task uncertainty and complexity. Our study has two important implications. First, it provides the theoretical groundwork for closely linking unique characteristics of the distinct brain regions in the context of RL. Second, our implementation is performed in a simple matrix form that accommodates expansion into biologically-plausible, highly-scalable, and generalizable neural architectures.

## 1. Introduction

One of the key challenges of reinforcement learning (RL) is to resolve environmental uncertainty and complexity with limited resources. In spite of the recent developments in machine learning to deal with complex problems [1–9], RL algorithms often have a hard time learning simple tasks that efficiently learnt by animals [10].

One common criticism is that a majority of learning algorithms based on deep learning have little biological relevance, though one can find some commonalities at the conceptual level [8, 9, 11–14]. Furthermore, our understanding of how the biological system implements learning is limited to simple tasks. A few working theories successfully explain the biological principles of each part of the brain, but they do not yet account for the ways in which interactions between different brain regions lead to optimal learning of the environment. Another limitation is that biological models for investigating high-level brain functions, such as context or memory, are not tightly linked with biological processes at the single-neuron level or accompany many biological constraints that limit their applicability.

To reconcile the discrepancies between RL algorithms and biological RL, here we explore biological processes that potentially contribute to efficient learning. Based on recent neural findings explaining the ability of the hippocampus and prefrontal cortex to efficiently glean space and time information from the environment, we propose a novel RL model to reliably learn a highly uncertain environment. This paper is structured as follows.

- First, we propose a novel spacetime encoding method by combining successor representation with theta phase precession. This module acts as a rollout prediction.
- Second, we implement prefrontal population coding for reliable reward prediction, called Q switch ensemble, that exploits the spacetime information of the episodes from the first module.
- Third, to further improve learning efficiency, we propose a single biological learning rule, accommodating both hippocampal-prefrontal replay and synaptic homeostasis. This learning rule subserves confidence-based metacognitive learning.
- Finally, we designed a task array simulating different levels of environmental uncertainty and complexity to demonstrate the capacity of our model, called spacetime Q-Network (STQN). We also ran systematical ablation analyses to show the unique contributions of each component to resolving task uncertainty and complexity.

## 2. Related Works and Neural Basis

The successor representation (SR) is a concept that represents the temporal proximity between the current, past, and future state of the environment [15]. As it captures the proximity of two events being relational, it has become a valuable tool in various fields, such as human reinforcement learning [16], robotic control [17], and visual navigation [18]. Recently, SR has gained significant attention in deep reinforcement learning [19, 20]. However, it has so far been used as a subordinate tool; for example as the realization of the posterior sampling [21], the utility for feature representation with a set of expectations [22], or the intermediate for inference using a nonparametric Dirichlet process mixture model [23]. In this regard, they have limited biological relevance.

Next, the hippocampal replay, specifically when awake, is the high-frequency neural activity inside the envelope of a sharp wave ripple (SWR) that occurs during resting or receiving a reward. SWR has long been considered as the bridge between short- and long-term memory consolidation [24]. There are several studies on this relationship; however, most of them have focused on relating the generative and terminative mechanism of the hippocampal replay to various synaptic properties [25, 26]. One study attempted to interpret the replay as iterative executions of Bellman backup and showed that their model can imitate some basic properties of the hippocampal replay [27]. However, this study neither examines the correspondence between their model and the prefrontal-hippocampal network nor explains the biological mechanisms of initiating, proceeding, and terminating replay.

Third, the synaptic homeostasis is a consistent synaptic operation needed to maintain the stability of neural networks undergoing rapid spike timing dependent plasticity (STDP), to properly store or process information [28]. This demand is because of the exponential property of the STDP network; the amounts of weight updates depend on the weight itself. This characteristic sometimes leads to destructive forgetting, masking of useful information the agent had learned, under the pathological conditions [29]. While a few studies have investigated the effect of synaptic homeostasis on learning [30–32], none of them are sufficient to carry real-time regulations during complex learning.

## 3. Methods

### 3.1. Spacetime Encoding

To build an effective encoder, we combined two concepts; successor representation (SR) and theta phase precession. To model the spatial properties of the place cell, successor representation is a rational choice as it can explain the irregular modulations of its receptive field [33] and be simulated by the spiking neural network [34]. Moreover, recent observations show that place cells can represent locations of the other animals enable the SR matrix to encode the target object [35, 36].

Next, we assume that the theta phase precession sharpens the spatial coding by adding temporal context to the place cell firing sequences. This hypothesis is reinforced by various experiments. For instance, it is observed that the hippocampal theta sequence tends to proceed toward the current goals [37]. The theta phase can divide the trajectory that an animal had passed and will pass into temporal bins [38] support this idea. Moreover, our assumption is assisted by an experiment that shows that pharmacological inhibitions of the lateral septum (LS), the major source of hippocampal theta wave, extinguish the ability to navigate without disturbing the intact place cell activities [39] or an experiment that shows that LS extends the egocentric firing of place cells to the entire environment located by the animal [40], which is an invaluable tool for planning and tackling further situations inside the complex spatial learning [41].

#### 3.3.1. Successor Representation for Predictive Coding

Assume that there are *N*_*H*_ place cells *c*_*H*_, which have the center of their receptive fields 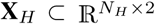. To represent the correlational relationship between the place cells, we used SR to represent the correlational relationship between the place cells. The SR matrix ***M*** encodes the expected and discounted future occupancy of the cell *c′*_*H*_ along a trajectory starting from the cell *c*_*H*_:

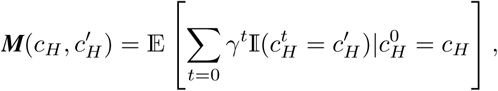

where 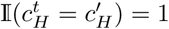 if 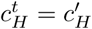, otherwise 0 [15]. To update the SR matrix, we group the place cells whose receptive fields contain an external stimuli (e.g., a ball in Pong game) inside. Specifically, if the Euclidean distance between the center **x**_*H*_ of the place cell *c*_*H*_ and the location of the input stimulus 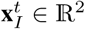 is smaller than the update distance *l*_*U*_, the cell *c*_*H*_ is in the candidate set,

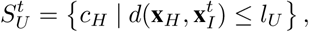

If 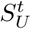 and 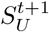 are not empty, the update begins. For every cell pairs 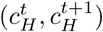 from 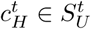 and 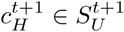, we do TD online learning for those pairs [42]:

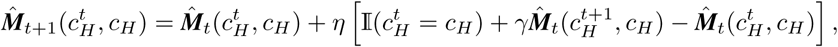

where *η* is a learning rate, *γ* is a discount factor, respectively.

#### 3.1.2. Theta Phase Precession for Spacetime Encoding

It is known that the theta phases inside the CA1 and PFC align in a regular manners [43]. To imitate this systematic modulation, we divide the single period of theta wave, between two peaks, into four equal parts. Then, we allocate the first, second, and third parts, respectively, to the past, current, and future place cells individually. The firing of PFC cells locates at the peak behind; therefore, the firing sequences of future cells become closest among the place cells. The sign of the cosine value is the criteria used to divide the past and future cells among all fired place cells. Assume that the input stimulus is located at **x**_*I*_ with a velocity of 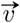. After defining the vector from the center **x**_*H*_ of place cell *c*_*H*_ to **x**_*I*_, we can discern whether or not the moving stimulus had already passed the center **x**_*H*_.

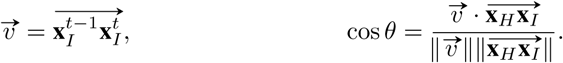

If the cosine value is negative, we can state that the cell *c*_*H*_ lies on the past trajectory of the input stimulus and vice versa. However, the dot product above can only separate past and future cells, not current cells. To compensate for this limitation, we define the current cell as cells closer to the input stimulus than the others. Mathematically, a set of current cells is:

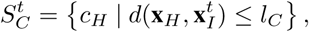

where *l*_*C*_ is a new parameter, the current distance.

#### 3.1.3. SR Firing Rate for Online Learning

The agent needs a channel to transfer processed spatiotemporal information. One of the biologically-plausible candidates is the firing sequence from the presynaptic to the postsynaptic neurons, which is supported by the LS [44]. However, based on our knowledge, there are no trials or attempts to use SR as the basis for modeling real-time activity of place cells during complex learning, specifically, not as an alternative representation of the abstract “state.” Therefore, we defined the firing rate for the SR matrix in online learning. From a set of current place cells, we define the firing rates of individual place cells. One way is to average the column vectors of current place cells in the SR matrix ***M***:

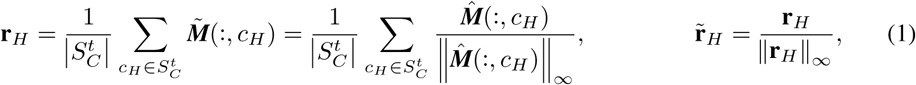

where 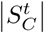 is the number of current place cells, 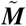 is normalized SR matrix obtained by dividing each column of the learned SR matrix by its maximum element. After comparing the rate vector **r**_*H*_ to the uniform random vector, we build **f**_*H*_, binary firing vector of all place cells.

### 3.2. Q Switch Ensemble

Q switch ensemble consists of two components: PFC cells and their population coding for reward prediction. In this study, our PFC cell model is based on the experiment that shows showing that individual PFC cells represent the difference between Q values of two options (left or right) to its firing rate while solving the Markov decision problem [45]. Based on this observation, we generalize PFC cell activity to binary Q classifiers, continuously updating its Q value and firing when it predicts a specific event will happen, of which firing rate is the function of Q value. Based on the recent evidence on dopaminergic neurons having different criteria of reward feedback [46] and the capability of PFC cells to encode spatial information [43], we represent the reward prediction in the form of population coding. Similar to the experimental data [43], each PFC cells inherit only the least spatial information, acting as a decision center. The PFC cells fire when it forecasts that the reward event will occur above the center of decision, then the agent compiles the votes to predict the location.

#### 3.2.1. Phase Distance for an Effective Weight Representations

Based on our assumption of the theta phase precession, there exists a relative attenuation of the neuronal signal intensity of the early (past) spikes, when compared to the late (future) spikes. Therefore, we calculated the effective coefficients following cable theory [47] and spike timing dependent plasticity (STDP) [48]. Specifically, using the equal phase distances between past-current, current-future, and future-PFC, we proposed the following weighting scheme:

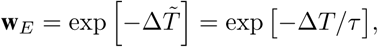

where 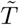 is a time distance divided by time constant *τ*, to convert the exponents to the integer.

#### 3.2.2. PFC Cell Population Coding for Binary Q Classification

To use the transferred temporal contexts by the phase distance, we multiplied the binary firing vector **f**_*H*_ with the effective weight **w**_*E*_, element-wisely, producing effective firing vector 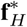. The PFC cells receive and combine 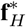 linearly through the weight matrix 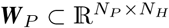, making the Q values,

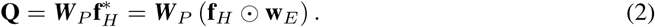

The firing rate of each PFC cell is a sigmoid function of the Q value. To separate the activity of PFC cells maximally, we used the mid-range of **Q** as the offset. To compute the firing rate of PFC cells **r**_*P*_:

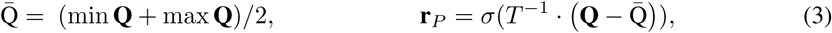

where *T*^−1^ is the inverse temperature of the sigmoid function *σ*.

#### 3.2.3 PFC Cell Population Coding for Reward Prediction

We interpreted the learning as a population process, starting from the binary Q classifier. We assume that there are *N*_*P*_ PFC cells *c*_*P*_, and their center of decision is 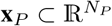. If the PFC cell predicts that the reward event will occur above its center of decision, it fires. Otherwise, when it forecasts that the reward will occur below its center, it does not fire. We then define the level of confidence, which quantifies the amount by which the single cell is confident with its decision,

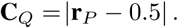

Here, the reward prediction is formalized in population coding. If the PFC centered at *x*_*P*_ fires, it indicates that the reward event was predicted to occur anywhere above the center of decision of the PFC cell. To compress the continuous domain of the prediction, we choose points arbitrarily for discretization. In our simulations, we select the midpoints between every decision centers and build a set 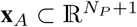. We refer to this set as the action set. The certainty **C**_*Q*_ now becomes a poll value. For a single action point *x*_*A*_, the firing of the single PFC cell centered at *x*_*P*_ provides the following:

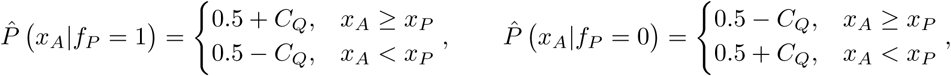

where *f*_*P*_ ∈ **f**_*P*_ is a Boolean variable whether or not this PFC cell mentioned above fired and *C*_*Q*_ ∈ **C**_*Q*_ is a corresponding individual confidence of the cell. After summing all the votes for the midpoints and then normalizing them, a discrete probability distribution over **x**_*A*_ can be obtained. The coordinate 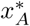 becomes the target; if the agent locates below 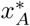, the agent moves up and so on.

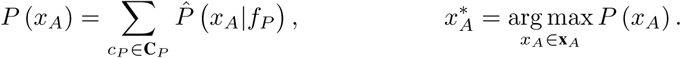

### 3.3. Learning with HPC-PFC Replay

In this study, we view the HPC-PFC replay as a recursive TD learning process. First, we establish a hypothesis reasoned from optogenetic manipulation, in which the sharp wave ripples modulate the synaptic weights [49]. Furthermore, this view is supported by the evidence that the relative progress of task learning systemically modulates the trend of replay [50], and it actively selects the highly rewarded event [51]. When the trial ends, the agent receives the reward information (e.g., the actual location at which the ball arrived). From this spatial information, every PFC cell receives its distinct rewards via the dopaminergic pathway [52]. Then, the learning process is initiated and proceeds recursively, underpinned by recurrent, interlamellar connections inside the hippocampus [53].

When the action point *x*_*B*_ is the closest to the one associated with reward (e.g., the location at which the ball arrived), the initial feedback signal for learning is simply the logical result of comparison; whether or not the point ball arrived was above the decision center of each PFC cell:

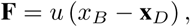

where *u* (⋅) is the unit step function. Then, the changes of synaptic weights are linear to the difference between the feedback and the firing rate of each PFC cell [42]. Moreover, to present the mechanism by which various phenomena seemingly contribute to the learning, we designed two simple functions; H (⋅) is the entropy learning rate function, *γ* (⋅) is the synaptic stability function (The details are in the supplementary material). With these utilities, the equation becomes:

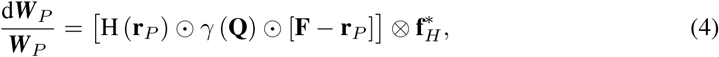

where 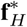 is the effective firing vector. After the learning is initiated, the agent must prepare the next interactions not just a right in front, but from a few steps behind. We assume that the HPC-PFC network achieves this by relaying the Q values and attempting to narrow the gap between adjacent decisions via weight modulation. Following the statistics and properties of the hippocampal replay, we implement the recursive algorithm to simulate such a behavior (supplementary material).

## 4 Simulation Result

### 4.1. Architecture of STQN

Our model consists of two components. First, the spacetime SR models the spatiotemporal encoding of the hippocampus by combining the successor representation as the spatial processor with the theta phase precession as the temporal channel. This module receives the current coordinate of the external input **x***I* and outputs the firing sequence of the relevant place cells. Next, it integrates the temporal order by multiplying the effective weight element-wise for the theta phase precession. Second, the Q switch ensemble receives the processed firing sequence and outputs the most probable reward location. After receiving the reward, the recursive TD learning updates the weights ***W***_*P*_, with respect to the difference between the decision certainty of each PFC cell and the relative feedback signal.

### 4.2. Novel task design varying the degree of uncertainty and difficulty level

We designed our task based on the classic Atari game Pong. To incorporate both the structural complexity and state-transition uncertainty into the task, we added three walls to test the learning performance of various agents. First, we doubled the width and height of the Pong table, four times larger than the OpenAI gym Pong [54], without modifying the parameters of the agent behavior. Next, we varied the amount of uncertainty in ball bouncing, by making the trajectory of the ball difficult to predict. After the ball bounces off the wall, a uniform noise is added to the angle of the velocity vector. This noise is bounded by the "variation angle" *θ*_*V*_. Using *θ*_*V*_, we can incorporate the uncertainty in a controllable, natural manner. Additionally, the environment is modulated in a dynamic manner, from regular oscillation to the chaotic Brownian motion. Lastly, we implemented the opponent with an ideal agent. For this, we assumed that the agent has complete access to the environmental structure (e.g., the exact location where the ball will arrive, given the current position and velocity vector of the ball). At every single frame, the opponent (ideal agent) receives the future location of arrival, but it contains additive white Gaussian noise. The task difficulty can be varied by manipulating the probability with which the opponent makes mistakes (standard deviation of this noise *σ*_*w*_). The difficulty level allows us to assess the capacity of the agent since the lesser mistakes the opponent makes, the longer the rally length becomes (Details are in supplementary material).

We set two hyperparameters, the "variation angle" *θ*_*V*_ and the standard deviation of opponent noise *σ*_*w*_. Subsequently, we built six tasks by combining two hyperparameter sets; 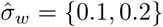, and *θ*_*V*_ = {0°, 10°, 30°} where 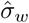 is the normalized standard deviation of the opponent, *σ*_*w*_ divided by the half-height of the Pong table. For simplicity, we hereinafter refer to the conditions 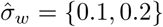 as EASY and HARD, and *θ*_*V*_ = {0°, 10°, 30°} as LOW, MID, and HIGH uncertainties, respectively.

### 4.3. Performance Comparison

To examine the way in which various deep RL agents deal with the aforementioned dynamic environments, we chose DQN [55], IQN [56], PPO [57], A3C-LSTM [12] as the representative comparison models. Notably, we found that our model is the only one that achieved guaranteed performances over the broad spectrum of uncertainty and complexity (Table 1) without the requirement of fine-tuning.

**Table 1:** Benchmark test (5M episodes)

| Uncertainty | Score |  |  |  |  |  | Rally length (100 Frames) |  |  |  |  |  |
| --- | --- | --- | --- | --- | --- | --- | --- | --- | --- | --- | --- | --- |
|  | LOW |  | MID |  | HIGH |  | LOW |  | MID |  | HIGH |  |
|  | EASY | HARD | EASY | HARD | EASY | HARD | EASY | HARD | EASY | HARD | EASY | HARD |
| DQN | -11.2 | -10.5 | -10.7 | -8.09 | -14.1 | -18.1 | 0.74 | 1.01 | 0.76 | 1.23 | 0.80 | 0.80 |
| IQN | -14.9 | -15.8 | -15.7 | -17.64 | -14.0 | -16.7 | 0.86 | 1.06 | 0.84 | 0.91 | 1.01 | 1.04 |
| PPO | -18.5 | -19.2 | -18.6 | -19.51 | -18.0 | -19.2 | 0.61 | 0.62 | 0.61 | 0.62 | 0.68 | 0.69 |
| A3C-LSTM | -9.4 | -14.2 | -10.0 | -14.66 | -11.0 | -15.1 | 1.15 | 1.33 | 1.18 | 1.35 | 1.17 | 1.30 |
| STQN* | **15.2** | **10.2** | **14.1** | **9.1** | **14.1** | **9.2** | **3.99** | **6.36** | **4.51** | **6.59** | **4.44** | **6.84** |

There are a few key points to note in RL learning. First, the score tended to decrease as the task difficulty increased, except for DQN. This is because DQN is unstable and it generally failed, but the minor (<15%) succeed to learn (supplementary material). Second, PPO failed to learn completely and this deficiency correlates significantly to task difficulty (supplementary material). The third is the reliability of the meta-RL agent (A3C-LSTM). To test its long-term learning, we increased the training episodes to 20M; however, we found that the effects are marginal (supplementary material). We used the rally length to quantify the capacity of the RL agents (see Table 1). When the capacity of the learning agent was comparable to that of the ideal agent (opponent), the rally length was observed increase abruptly, since there were no mistakes to terminate the rally. We were able to identify this effect in our model (>100%), regardless of the task uncertainty. It clearly demonstrated the high capacity of our model and also implied that the policy of our model is near-optimal with the scoreboard. However, this effect was not very pronounced in the other models (<20%) and tended to disappear under high uncertainty conditions, thereby indicating incomplete learning in these models.

### 4.4. Role of Predictive Spacetime Encoding in Learning (Ablation Study)

To examine the unique roles of individual components, we conducted the ablation study. More details on analyses and discussions are provided in Supplementary material. First, we quantified the degree of contribution of theta phase precession on task performance. Next, we examined how place cell encodes information and supports complex prediction by modulating the two hyperparameters; the update distance *l*_*U*_ and the current distance *l*_*C*_ based on successor representation. In here, we introduce two different measures, the "error" *E* and the "accuracy" *RC* (see Figure 3). In-depth performance analysis and discussions about biological implications are provided in the supplementary material.

#### 4.4.1. Theta Phase Precession

The theta phase precession implements the idea that the temporal distance controls the relative contribution of a single spike to the remaining process. This modulation allows the agent to take a predictive "vector decision": one should act differently if the situation were predicted to change in the foreseeable future. As learning proceeds, the learning agent becomes sensitive to the environmental changes by encoding the "context". However, the effectiveness of this strategy is constrained by the task uncertainty and the task difficulty.

By comparing the time courses of learning of the agent with and without theta phase precision, we found its significant contribution on performance (figure 2). We also found that the terminal error difference depends on the uncertainty and the difficulty level of the task. Especially, the effect of task uncertainty on the performance wears off in the high task difficulty conditions (Hard).

**Figure 1:**
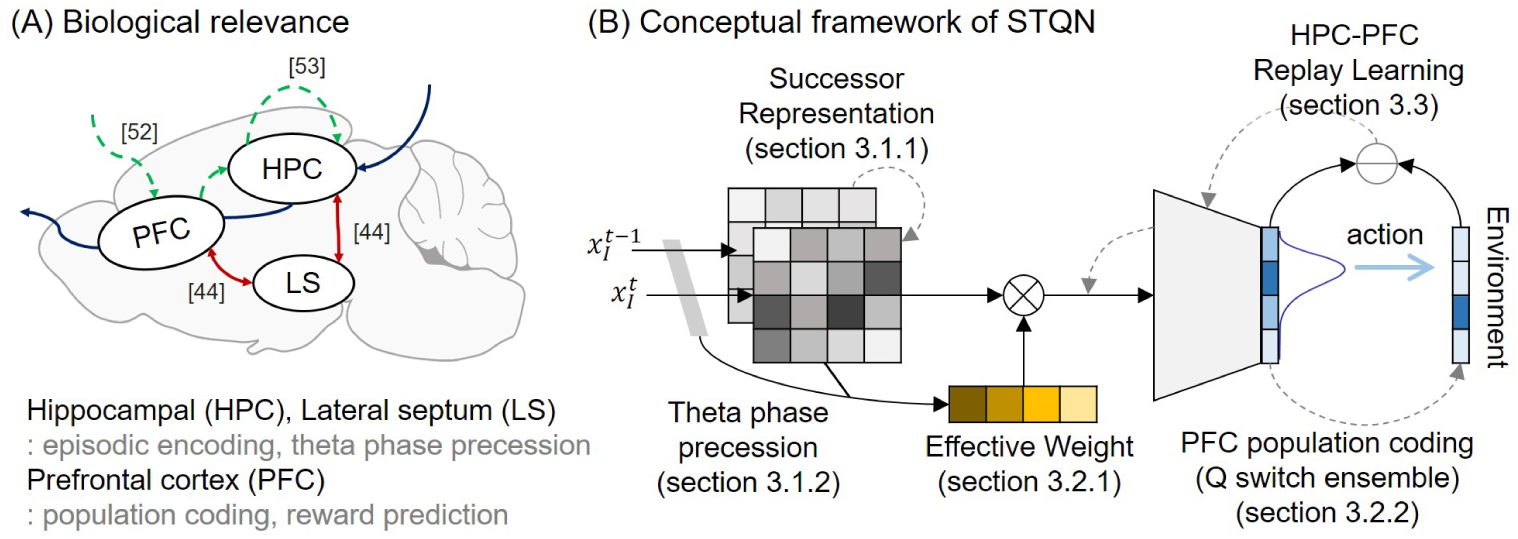
Concept Diagram of STQN

**Figure 2:**
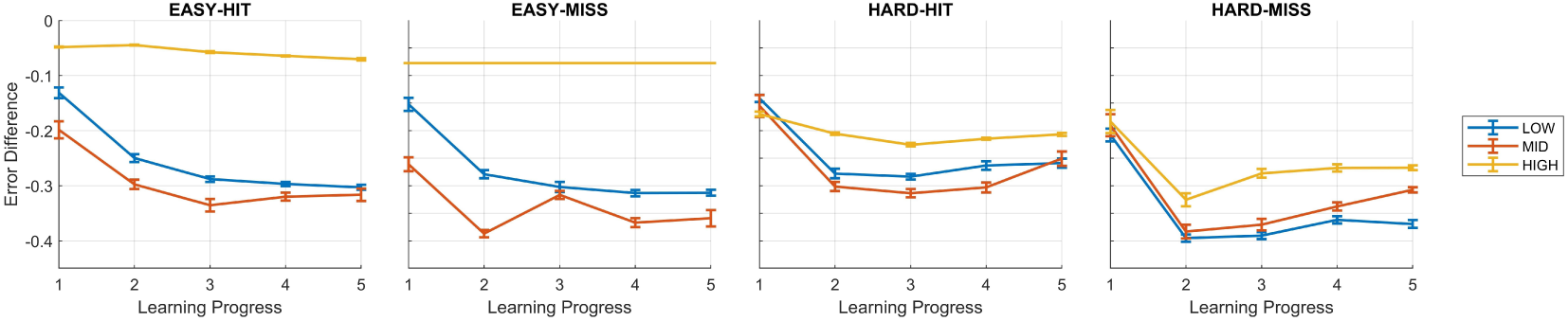
Theta Phase Precession HIT corresponds to the case when the agent successfully passes the ball MISS corresponds to the case when the agent makes mistakes and receives negative reward

#### 4.4.2. Successor Representation

Update distance *l*_*U*_ is the hyperparameter that represents the number of future states that a single current state should encode. The higher the uncertainty, the harder it is to predict future states from the given current state; consequently, the time required to achieve a certain level of accuracy also increases. This leads to the two distinct properties; (1) the slope of the error with respect to *l*_*U*_ becomes more negative proportionally (Figure 3A Amplitude), and (2) the error difference between the agent with the normal *l*_*U*_ and the small value increases (Figure 3A Offset) with the decrease in the task uncertainty.

**Figure 3:**
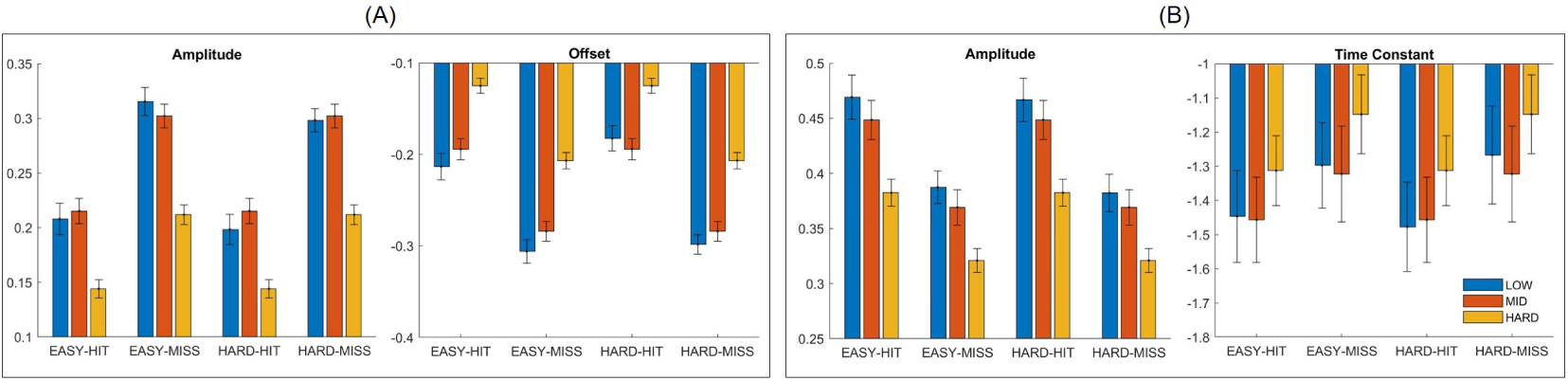
(A) Update Distance and (B) Current Distance Fitting with (A) 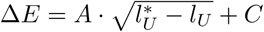 and (B) *R*_*C*_ = *A* ⋅ exp[*τ*_*C*_ ⋅ *l*_*C*_]

If we view the SR matrix as an encoder and the Q switch ensemble as a decoder, then the current distance *l*_*C*_ is translated into the factor modulating the amount of information that the encoder can process. This provides two predictions; (P1) the agent needs more capacity in highly uncertain environment, and (P2) the measure *R*_*C*_, the ratio of "correct" PFC cells, decreases when the amount of uncertainty increases with the same *l*_*C*_. According to (P1), the difference of *R*_*C*_ between the agents with the changed and normal *l*_*C*_, ∆*R*_*C*_, should decrease more rapidly (Figure 3B Time Constant) since 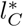 that makes ∆*R*_*C*_ =0 decreases, where 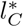 is the minimal *l*_*C*_ that can support sufficient number of symbols (lower bound of *l*_*C*_). According to (P2), the value at infinitesimal *l*_*C*_ increases when the amount of task uncertainty decreases (Figure 3B Amplitude).

## 5 Summary and discussion

Motivated by recent neural findings about space-time information processing in the hippocampus and prefrontal cortex, we propose a novel spacetime RL framework, called spacetime Q-Network (STQN), that reliably learns highly uncertain environment. The implications of our study are as follows. First, our spacetime encoding allows us to examine the hypothesis that hippocampal place cells with theta phase precession facilitate predictive encoding and learning. Second, we demonstrate that the group of binary Q classifiers as a proxy for PFC cells can predict the complex physical processes [46]. Third, the proposed learning rule underlines a direct relevance of the homeostatic synaptic plasticity to Q-learning [45]. In summary, our study lays the theoretical groundwork for integrating unique properties of separate brain regions in the context of RL. In addition, our formulation of biological processes in the form of a simple matrix offers valuable insights into biologically-plausible and highly-scalable neural architecture designs.

### Broader Impact

This work is based on the computer simulation, so there is no direct impact on animals or humans. Therefore, there are no potential ethical issues. This work has the following potential impacts on society: The design of novel deep learning architecture. We proposed a biologically plausible RL model integrating unique characteristics of distinct brain regions. With taking into account a brain’s capacity encoding the space and time information, the proposed model exploits the reliable learning the highly uncertain and complex environment. The model was implemented in a simple matrix form, so it is expected to accommodate expansion into highly-scalable and generalizable neural architecture. The training process is based on simulations in the virtual (game) environment. Hence, there is no psychological or physical harm to humans inflicted by our model.

## Supporting information

example task video

