## supplementary material for "Biological Reinforcement Learning via Predictive Spacetime Encoding"

**Minsu Abel Yang**
KAIST
Department of Bio and Brain Engineering
Department of Electrical Engineering


**Jee Hang Lee**
Sangmyung University
Department of Human-Centered AI


**Sang Wan Lee**
KAIST
Department of Bio and Brain Engineering


**3 Environment Specifications** **9**

Preprint. Under review.

**4 Analysis 1 - Task Performance** 9
4.1 Definition of Performance Measures 9
4.1.1 Score 9
4.1.2 Rally Length 9
4.2 Experimental Procedure 9
4.2.1 Performance Comparison 9
4.2.2 Ablation Study 10
4.3 Results 11
4.3.1 Performance Comparison 11
4.3.2 Ablation Study 14

**5 Analysis 2 - Biological Implication** 16
5.1 Additional Performance Measures for In-depth Analyses 16
5.1.1 Relative Time 16
5.1.2 Timing Error 16
5.1.3 Cell Population Accuracy 16
5.2 Experimental Procedure 17
5.2.1 Theta Phase Precession 17
5.2.2 Successor Representation - Update Distance 17
5.2.3 Successor Representation - Current Distance 17

**6 Role of Predictive Spacetime Encoding in Learning** 18
6.1 Theta Phase Precession 18
6.2 Successor Representation 18
6.2.1 Update Distance 18
6.2.2 Current Distance 20

**Notation 1:** Input

| | |
|---|---|
| $\mathbf{x}_I^t$ | Input stimulus at time $t$ |
| $\mathbf{d}_I^t$ | The distance vector at time $t$ |
| $\overrightarrow{v}$ | The velocity vector of the input |

**Notation 2:** Place Cell

| | |
|---|---|
| $c_H$ | Place cell |
| $d_L$ | The unit lattice length |
| $N_H$ | The number of place cells |
| $\mathbf{X}_H$ | The center of place cell receptive fields |
| $l_U$ | Update distance |
| $\eta$ | Learning rate of SR matrix update |
| $\gamma$ | Discount factor of SR matrix update |
| $\mathbf{b}_U^t$ | The inclusion indicator of the set $S_U^t$ |
| $S_U^t$ | A set contains place cells target of update at time $t$ |
| $\hat{\boldsymbol{M}}_t$ | SR matrix at time $t$ updated by online learning |
| $\tilde{\boldsymbol{M}}_t$ | Normalized SR matrix at time $t$ |
| $l_C$ | Current distance |
| $\mathbf{b}_C^t$ | The inclusion indicator of the set $S_C^t$ |
| $S_C^t$ | A set contains current firing place cells at time $t$ |
| $\mathbf{r}_H$ | The firing rate of entire place cells |
| $\hat{\mathbf{r}}_H$ | The normalized firing rate of entire place cells |
| $\mathbf{f}_H$ | The firing vector of entire place cells |
| $\mathbf{w}_E$ | The effective weight by the theta phase precession |
| $\mathbf{f}_H^*$ | The effective firing vector |

**Notation 3:** PFC Cells

| | |
|---|---|
| $c_P$ | PFC cell |
| $N_P$ | The number of PFC cells |
| $\mathbf{x}_P$ | The decision center of PFC cells |
| $\boldsymbol{W}_P$ | HPC-PFC weight matrix |
| $\mathbf{Q}$ | The Q value of entire PFC cells |
| $T^{-1}$ | The inverse temperature |
| $\mathbf{r}_P$ | The firing rate of entire PFC cells |
| $\mathbf{f}_P$ | The firing vector of entire PFC cells |
| $\mathbf{C}_Q$ | The confidence level of entire PFC cells |

**Notation 4:** Prediction

| | |
|---|---|
| $\mathbf{x}_A$ | A set contains possible position of the agent |
| $P(\cdot)$ | The predictive probability distribution over $\mathbf{x}_A$ |
| $x_A^*$ | The arguments of the maxima of $P(\mathbf{x}_A)$ |

**Notation 5:** Replay

| | |
|---|---|
| $e_M$ | The maximum of entropy learning rate |
| $\mathbf{F}$ | The relative dopaminergic feedback signal |
| $H(\cdot)$ | The entropy learning rate function |
| $\gamma(\cdot)$ | The synaptic stability function |

### 1 Methods in Matrix Form

#### 1.1 Spacetime Encoding

##### 1.1.1 Successor Representation for Predicting Coding

Assume that there are $N_H$ place cells $c_H$ , which have the center of their receptive fields $\mathbf{X}_H \subset \mathbb{R}^{N_H \times 2}$. To represent the correlational relationship among the place cells, we use SR to represent $\boldsymbol{M}$. To update the SR matrix, we group the place cells whose receptive fields contain an external stimulus (e.g., a ball in Pong game) inside. We denote the Euclidean distance between the center of the place cells as $\mathbf{X}_H$ and the location of the input stimulus as $\mathbf{x}_I^t \in \mathbb{R}^{1\times 2}$ to $\mathbf{d}_I^t$:

$$\mathbf{d}_I^t = \mathrm{sqrt}\left(\mathrm{diag}\left(\left(\mathbf{X}_H - \mathbf{x}_I^t\right)\cdot\left(\mathbf{X}_H - \mathbf{x}_I^t\right)^T\right)\right)$$

where $\mathrm{diag}(\cdot)$ is a function that returns main diagonals of the matrix inside as a column vector and $\mathrm{sqrt}(\cdot)$ is a function that returns the square root of the vector inside element-wise. Specifically, if an element of $\mathbf{d}_I^t$ is smaller than the update distance $l_U$, the corresponding cell $c_H$ is in the candidate set. Mathematically, for a candidate set $S_U^t$, its binary vector is

$$S_U^t = \left\{c_H \mid d(\mathbf{x}_H, \mathbf{x}_I^t) \le l_U\right\}, \qquad \mathbf{b}_U^t = u\left\{l_U \mathbb{1} - \mathbf{d}_I^t\right\},$$

where the element of binary set vector $\mathbf{b}_U^t$ becomes 1 if and only if the corresponding place cell $c_H \in S_U^t$. Here, $\mathbb{1}$ is a column vector that contains only 1, and $u(\cdot)$ is the unit step function.

If $S_U^t$ and $S_U^{t+1}$ are not empty, the update begins. For every cell pair $(c_H^t, c_H^{t+1})$ from $c_H^t \in S_U^t$ and $c_H^{t+1} \in S_U^{t+1}$, we perform TD online learning [1]. Using $b_U^t$ and $b_U^{t+1}$, the update rule, matrix form, is

$$\hat{\boldsymbol{M}}_{t+1}(\mathbf{b}_U^t,:) = \hat{\boldsymbol{M}}_t(\mathbf{b}_U^t,:) + \eta\left[\boldsymbol{I}_{N_H}(\mathbf{b}_U^t,:) + \gamma\hat{\boldsymbol{M}}_t(\mathbf{b}_U^{t+1},:) - \hat{\boldsymbol{M}}_t(\mathbf{b}_U^t,:)\right],$$

where $\boldsymbol{I}_{N_H} \subset \mathbb{R}^{N_H \times N_H}$ is an identity matrix, $\eta$ is the learning rate, and $\gamma$ is the discount factor.

##### 1.1.2 Theta Phase Precession for Spacetime Encoding

As mentioned in the main paper, we divide the single period of the theta wave between two peaks into four equal parts. Then, we allocate the first, second, and third parts individually to the past, current, and future place cells, respectively. The firing of PFC cells locates at the late peak among two peaks. Moreover, the sign of the cosine value is the criteria used to divide the past and future cells among all fired place cells. Assume that the input stimulus is located at $\mathbf{x}_I$ with a velocity of $\vec{v}$. After defining the vectors from $\mathbf{x}_I$ to the center of place cells $\mathbf{X}_H$, we can discern whether the moving stimulus has already passed the center $\mathbf{x}_H \subset \mathbf{X}_H$:

$$\vec{v} = \overrightarrow{\mathbf{x}_I^{t-1}\mathbf{x}_I^t}, \qquad \mathbf{s}_T = \mathrm{sgn}\left(\cos\theta\right) = \mathrm{sgn}\left(\left[\left(\mathbf{X}_H - \mathbf{x}_I^t\right)\cdot\frac{\vec{v}^T}{\|\vec{v}\|}\right]\oslash\mathbf{d}_I^t\right).$$

where $\mathrm{sgn}\left(\cdot\right)$ is the sign function. If the cosine value is negative, we can state that cell $c_H$ lies on the past trajectory of the input stimulus and vice versa. However, the dot product above can only separate past and future cells, not current cells. To compensate for this limitation, we define the current cell as cells closer to the input stimulus than the others. Mathematically, a set of current cells and its binary vector are

$$S_C^t = \left\{c_H \mid d(\mathbf{x}_H, \mathbf{x}_I^t) \le l_C\right\}, \qquad \mathbf{b}_C^t = u\left\{l_C \mathbb{1} - \mathbf{d}_I^t\right\},$$

where $l_C$ is a new parameter, the current distance. The element of binary set vector $\mathbf{b}_C^t$ becomes 1 if and only if the corresponding place cell $c_H \in S_C^t$.

##### 1.1.3 SR Firing Rate for Online Learning

After normalizing each column of the learned SR matrix $\hat{\boldsymbol{M}}$ by dividing it by its maximum element, we calculate the firing rates of individual place cells by averaging the column vectors of current place cells in the normalized SR matrix $\tilde{\boldsymbol{M}}$. After comparing the normalized rate vector $\tilde{\mathbf{r}}_H$ to the uniform random vector, we build $\mathbf{f}_H$, the binary firing vector of all place cells.

$$\tilde{\boldsymbol{M}} = \frac{\hat{\boldsymbol{M}}}{\|\hat{\boldsymbol{M}}\|_\infty}, \qquad \mathbf{r}_H = \frac{\tilde{\boldsymbol{M}} \cdot \mathbf{b}_C^t}{\sum \mathbf{b}_C^t} \qquad \tilde{\mathbf{r}}_H = \frac{\mathbf{r}_H}{\|\mathbf{r}_H\|_\infty}, \qquad \mathbf{f}_H = \mathcal{U}(0,1) \leq \tilde{\mathbf{r}}_H \tag{1}$$

#### 1.2 Q Switch Ensemble

##### 1.2.1 Phase Distance for an Effective Weight Representations

Based on our assumption of the theta phase precession, there exists a relative attenuation of the neuronal signal intensity of the early (past) spikes when compared to the late (future) spikes. Therefore, we calculated the effective coefficients following cable theory [2] and spike timing dependent plasticity (STDP) [3]. Specifically, using the equal phase distances between past-current, current-future, and future-PFC, we proposed the following weighting scheme:

**Table S1:** Effective Weight Setting

| Label | $\mathbf{f}_H$ | $\mathbf{s}_T$ | $\mathbf{b}_C^t$ | $\mathbf{w}_E$ |
|---|---|---|---|---|
| None | 0 | ... | ... | ... |
| Past | 1 | $-1$ | 0 | $1/e^3$ |
| Current | 1 | ... | 1 | $1/e^2$ |
| Future | 1 | $+1$ | 0 | $1/e^1$ |

The mathematical description of the effective weight satisfying the above conditions is

$$\mathbf{w}_E = \exp\left[\mathbf{s}_T \odot \left(1 - \mathbf{b}_C^t\right) - 2\right].$$

##### 1.2.2 PFC Cell Population Coding for Binary Q Classification

To use the transferred temporal contexts with the phase distance, we multiplied the binary firing vector $\mathbf{f}_H$ with the effective weight $\mathbf{w}_E$ element-wise, producing effective firing vector $\mathbf{f}_H^*$. The PFC cells receive and combine $\mathbf{f}_H^*$ linearly through the weight matrix $\boldsymbol{W}_P \subset \mathbb{R}^{N_P \times N_H}$, yielding the Q values

$$\mathbf{Q} = \boldsymbol{W}_P \mathbf{f}_H^* = \boldsymbol{W}_P \left(\mathbf{f}_H \odot \mathbf{w}_E\right). \tag{2}$$

The firing rate of each PFC cell is a sigmoid function of the Q value. To separate the activity of PFC cells maximally, we used the mid-range of $\mathbf{Q}$ as the offset. To compute the firing rate of PFC cells $\mathbf{r}_P$ and their corresponding binary firing vector $\mathbf{f}_P$:

$$\bar{\mathrm{Q}} = (\min \mathbf{Q} + \max \mathbf{Q})/2, \qquad \mathbf{r}_P = \sigma(T^{-1} \cdot (\mathbf{Q} - \bar{\mathrm{Q}})), \qquad \mathbf{f}_P = \mathcal{U}(0,1) \leq \mathbf{r}_P, \tag{3}$$

where $T^{-1}$ is the inverse temperature of the sigmoid function $\sigma$.

##### 1.2.3 PFC Cell Population Coding for Reward Prediction

We interpreted the learning as a population process, starting from the binary Q classifier. We assume that there are $N_P$ PFC cells $c_P$, and their center of decision is $\mathbf{x}_P \subset \mathbb{R}^{N_P}$. If the PFC cell predicts that the reward event will occur above its center of decision, it fires. Otherwise, if it forecasts that the reward will occur below its center, it does not fire. We then define the level of confidence, which quantifies the amount by which the single cell is confident with its decision:

$$\mathbf{C}_Q = |\mathbf{r}_P - 0.5| .$$

Here, the reward prediction is formalized in population coding. If the PFC centered at $x_P$ fires, it indicates that the reward event was predicted to occur anywhere above the center of decision of the PFC cell. To compress the continuous domain of the prediction, we choose points arbitrarily for discretization. In our simulations, we select the midpoints between every decision center and build a set $\mathbf{x}_A \subset \mathbb{R}^{N_P+1}$. We refer to this set as the action set. The certainty $\mathbf{C}_Q$ now becomes a poll value. For the entire action points $\mathbf{x}_A$,

$$\hat{P}\left(\mathbf{x}_A|\mathbf{f}_P\right) = \text{sgn}\left(\mathbf{x}_A - \mathbf{x}_P^T\right) \cdot \text{diag}\left(\text{sgn}\left(\mathbf{f}_P - 0.5\right)\right),$$

where diag$(\cdot)$ is a function constructs a square diagonal matrix with the elements of vector inside on the main diagonal. After summing all the votes for the midpoints and then normalizing them, a discrete probability distribution over $\mathbf{x}_A$ can be obtained. The coordinate $x_A^*$ becomes the target; if the agent locates below $x_A^*$, the agent moves up and vice versa.

$$P\left(\mathbf{x}_A\right) \propto N_P/2 + \hat{P}\left(\mathbf{x}_A|\mathbf{f}_P\right) \cdot \mathbf{C}_Q, \qquad x_A^* = \underset{x_A \in \mathbf{x}_A}{\arg\max}\, P\left(x_A\right).$$

#### 1.3 Learning with HPC-PFC Replay

##### 1.3.1 HPC-PFC Replay for Recursive TD Learning

When the action point $x_B$ is the closest to the one associated with reward (e.g., the location at which the ball arrived), the initial feedback signal for learning is simply the logical result of comparison; whether the point at which ball arrives is above the decision center of each PFC cell:

$$\mathbf{F} = u\left(x_B - \mathbf{x}_P\right),$$

Then, the changes of synaptic weights are linear to the difference between the feedback and the firing rate of each PFC cell [1]. Moreover, to present the mechanism by which various phenomena seemingly contribute to the learning, we designed two simple functions; $\text{H}(\cdot)$ is the entropy learning rate function, and $\gamma(\cdot)$ is the synaptic stability function. With these utilities, the equation becomes

$$\frac{\text{d}\boldsymbol{W}_P}{\boldsymbol{W}_P} = \left[\text{H}\left(\mathbf{r}_P\right) \odot \gamma\left(\mathbf{Q}\right) \odot \left[\mathbf{F} - \mathbf{r}_P\right]\right] \otimes \mathbf{f}_H^*, \tag{4}$$

where $\mathbf{f}_H^*$ is the effective firing vector. We assume that the HPC-PFC network achieves this by relaying the Q values and attempting to narrow the gap between adjacent decisions via weight modulation. Following the statistics and properties of the hippocampal replay, we implement a recursive algorithm to simulate such a behavior. We explain this after presenting the utility functions.

##### 1.3.2 Synaptic Homeostasis for Stabilizing Weight Updates

To protect the STDP learning system [3] against weight explosion, we build a simple regulator function, inspired by the shared log-normal distributions of synaptic weights [4] and synaptic homeostasis. With this regulator function, the distribution of **Q** and its contributing weights follow a log-normal distribution because it is guided by the first gradient of the natural logarithm function.

$$\gamma\left(\mathbf{Q}\right) = \frac{\mathbf{1}}{\mathbf{dQ}} \ln\left(1 + \mathbf{Q}\right) = \frac{1}{1 + \mathbf{Q}}$$

##### 1.3.3 Entropy Learning Rate

In the cortical level, the human brain is known to dynamically allocate its learning rate, i.e. constant resources, to address uncertain decisions in order to resolve the causal relationship between the stimulus and the outcome [5]. Moreover, the study found that the hippocampus engages actively to perform "switching" between one-shot learning and incremental learning [5] [6]. To fully reflect this mechanism in our model, we designed the uncertainty dependent learning rate function. If we assume that the PFC cell follows the Bernoulli distribution $p = \mathbf{r}_P$, we can calculate the uncertainty in the individual cell using binomial entropy as follows.

$$\mathbf{H}(\mathbf{r}_P) \propto \mathbf{H}_B(\mathbf{r}_P) = -\mathbf{r}_P \odot \ln \mathbf{r}_P - (1 - \mathbf{r}_P) \odot \ln(1 - \mathbf{r}_P) \qquad \mathbf{H}(\mathbf{r}_P) = \frac{e_M}{\ln 2} \cdot \mathbf{H}_B(\mathbf{r}_P)$$

#### 1.4 Algorithm

**Algorithm 1** HPC-PFC Replay

**Input:** $\mathbf{f}_H$ : Binary firing vector of place cells at last
**Input:** $\mathbf{r}_P$ : Firing rates of entire PFC cells at last

1: **function** REPLAY( )
2: $\quad \mathbf{r}_P^{-1} \leftarrow \mathbf{r}_P, \mathbf{f}_H^{-1} \leftarrow \mathbf{f}_H$
3: $\quad$ **while** true **do**
4: $\quad\quad \mathbf{f}_H \leftarrow$ Results from Eq. 1 with $\mathbf{b}_C^t = \mathbf{f}_H^{-1}$
5: $\quad\quad \mathbf{f}_H^0 \leftarrow \mathbf{f}_H \wedge \neg \mathbf{f}_H^{-1}$
6: $\quad\quad$ **if** $\exists \mathbf{f}_H^0$ **then**
7: $\quad\quad\quad \mathbf{Q} \leftarrow$ Results from Eq. 2 with $\mathbf{f}_H^* = \mathbf{f}_H^0$
8: $\quad\quad\quad \mathbf{r}_P^0 \leftarrow$ Results from Eq. 3 with $\mathbf{Q}$
9: $\quad\quad\quad$ Update using Eq. 4 with $\mathbf{F} = \mathbf{r}_P^{-1}$ and $\mathbf{r}_P = \mathbf{r}_P^0$
10: $\quad\quad\quad \mathbf{r}_P^{-1} \leftarrow \mathbf{r}_P^0, \mathbf{f}_H^{-1} \leftarrow \mathbf{f}_H^0$
11: $\quad\quad$ **else**
12: $\quad\quad\quad$ BREAK - END REPLAY
13: $\quad\quad$ **end if**
14: $\quad$ **end while**
15: **end function**

The algorithm 1 receives two inputs when the interaction with the environment terminates and the positive or negative reward is given: the binary firing vector of entire place cells, and the firing rates of entire PFC cells. In line 4, we select the next firing place cells from the first input. This is due to the assumption that the hippocampal replay concurrently fires adjacent place cells at each time frame. In line 5, as the hippocampal replay is a high-frequency neural activity (>200Hz), we consider the refractory period by turning off the firing place cells if they also fired at last. In line 6, the algorithm validates whether any firing place cells exist in the current time frame. If there exist, the update procedures continue and the function substitutes the contents for the recursive learning.

### 2 Model Implementation

#### 2.1 Architecture of STQN

The positions of the place cells are structured by the hexagonal lattice, and the unit length of the lattice is $d_L$. In addition, we add white Gaussian noise to the positions for the randomness. All the parameters related to the distance are divided by $d_L$ for simplicity.

**Table S2:** STQN Specifications

| | |
|---|---|
| $d_L$ | 15 |
| $N_H$ | $\simeq$750 |
| $l_U/d_L$ | 1.4 |
| $\eta$ | 0.1 |
| $\gamma$ | 0.7 |
| $l_C/d_L$ | 1.8 |
| $N_P$ | 100 |
| $T^{-1}$ | 1 |
| $e_M$ | 1.7 |

The unit lattice length of the place cell $d_L$ was determined to match the receptive field size of the place cells (10-20 cm radius). The number of place cells $N_H$ is automatically determined by placing place cells in a hexagonal pattern. The $l_U$ and $l_C$ are the free parameters for SR, and their effects on task performance were thoroughly analyzed in this paper. The free parameters for RL, the learning rate $\eta$ and discount factor $\gamma$, were found to be not highly sensitive to task performance. The number of PFC cells $N_P$ was chosen arbitrarily (=100), and the inverse temperature parameter $T^{-1}$ was fixed at the default level (=1). The maximum entropy learning rate $e_M$ is the only hyperparameter of this model.

#### 2.2 Architecture of DQN

We used the DQN agent [7] in the class DQN of stable-baselines [8]. From the default option, the DQN class has double Q learning [9] and dueling extensions [10]. Each policy network is a multilayer perceptron (MLP) with 2 layers of 64 hidden units for each.

#### 2.3 Architecture of PPO

We used the PPO agent [11] in the class PPO1 of stable-baselines [8]. Each policy network is a multilayer perceptron (MLP) with 2 layers of 64 hidden units for each.

#### 2.4 Architecture of IQN

We implemented the IQN agent, following the structure listed in [12]. You can find the actual code for the model and entire simulations in `https://AAA.AAA`

#### 2.5 Architecture of A3C-LSTM

We implemented the A3C-LSTM agent, following the structure listed in [13]. You can find the actual code for the model and entire simulations in `https://AAA.AAA`

### 3 Environment Specifications

**Table S3:** Environment Specifications

| | |
|---|---|
| Half width of Pong table | 256 |
| Half height of Pong table | 128 |
| Location line of Pong panel | 10 |
| Half width of Pong panel | 14 |
| Velocity of Pong panel | 10 |
| Velocity of Pong ball | 7 |

Specifically, the location line of the Pong panel, or simply "pad line," refers to the distance from the wall to where the pad is fixed in the horizontal direction. Because our task needs only vertical movement, we fixed the horizontal position of the agent. For instance, if the center of the Pong table is at the origin, two pads will be at $(256-10,\cdots)$ and $(-256+10,\cdots)$, respectively.

### 4 Analysis 1 - Task Performance

#### 4.1 Definition of Performance Measures

##### 4.1.1 Score

The scoring rules are identical to table tennis. The left panel $x<0$ is for the opponent and the right panel $x>0$ is for the agent. The scoreboard is in the shape of tuple $(\mathrm{A}:\mathrm{B})$, A for the opponent and B for the agent. When the left panel misses the ball, the agent gets a point and B increases by 1, and vice versa. If either side reaches the end-score of 21, the game terminates. The definition of the measure "score" is $\mathrm{B}-\mathrm{A}$. Then, the range of the measure "score" is $[-21,21]$.

##### 4.1.2 Rally Length

Suppose that there are $L_R$ frames passed until the game terminates. Then, the definition of the "rally length" is the average length of a single rally, obtained by dividing $L_R$ by the number of rallies $\mathrm{A}+\mathrm{B}$.

#### 4.2 Experimental Procedure

The epochs are identical to 5M interactions (frames) for all simulations. As mentioned in the main paper, we tested all agents in 6 different tasks by varying the values of two hyperparameters $\theta_V$ and $\hat{\sigma}_w$. Subsequently, we combine two hyperparameter sets: $\hat{\sigma}_w=\{0.1,0.2\}$ and $\theta_V=\{0°,10°,30°\}$.

##### 4.2.1 Performance Comparison

We repeated all simulations 10 times to validate the learning ability of deep RL agents, except for DQN. For DQN, we repeated all simulations 30 times to investigate its instability and bifurcation. Moreover, to test the long-term learning ability of the meta-RL agent (A3C-LSTM), we extended the epochs to 20M and performed an additional 10 simulations. After simulation, we smoothed the raw data with simple exponential smoothing after the optimization and then discretized them into 11 bins of equal width. Moreover, to investigate the bifurcation of the DQN, we generated 10000 data points with 1D interpolation and the optimized exponential smoothing curve for each simulation. Then, we optimized the number of groups by using the silhouette score and clustered them.

###### 4.2.2 Ablation Study

To validate the contributions of the individual components of our model, we substituted or deleted the component and repeated all simulations 10 times. After simulation, we smoothed the raw data via simple exponential smoothing after the optimization and then discretized them into 11 bins of equal width. In addition, for the normalization of the score, we linearly transformed $[-21, 21]$ to $[0, 1]$. The details of the ablation study are following.

1. For the theta phase precession (TPP), we shuffled the non-zero elements of $\mathbf{w}_E$.
2. For the successor representation (SR), we replaced the SR matrix $\hat{\boldsymbol{M}}$ by a 2D Gaussian distribution. The firing rate of all place cells is

$$\mathbf{r}_H = \exp\left[-\frac{\mathbf{d}_I^t}{2 \cdot \sigma_G^2}\right], \qquad \tilde{\mathbf{r}}_H = \frac{\mathbf{r}_H}{\|\mathbf{r}_H\|_\infty}, \qquad \mathbf{f}_H = \mathcal{U}(0,1) \leq \tilde{\mathbf{r}}_H,$$

   where $\sigma_G$ is the standard deviation of the 2D Gaussian distribution to represent the radius of the place cell receptive field. We set $\sigma_G$ to 10.
3. For the synaptic stability function (SSF), we changed it to a constant, $\gamma(\mathbf{Q}) = 1$.
4. For the entropy learning rate (ELR), we changed it to a constant, the average value of $\mathbf{H}_B(\mathbf{r}_P)$.
5. For the zero-regulator case (XOR), we did both procedures for SSF and ELR.

#### 4.3 Results

##### 4.3.1 Performance Comparison

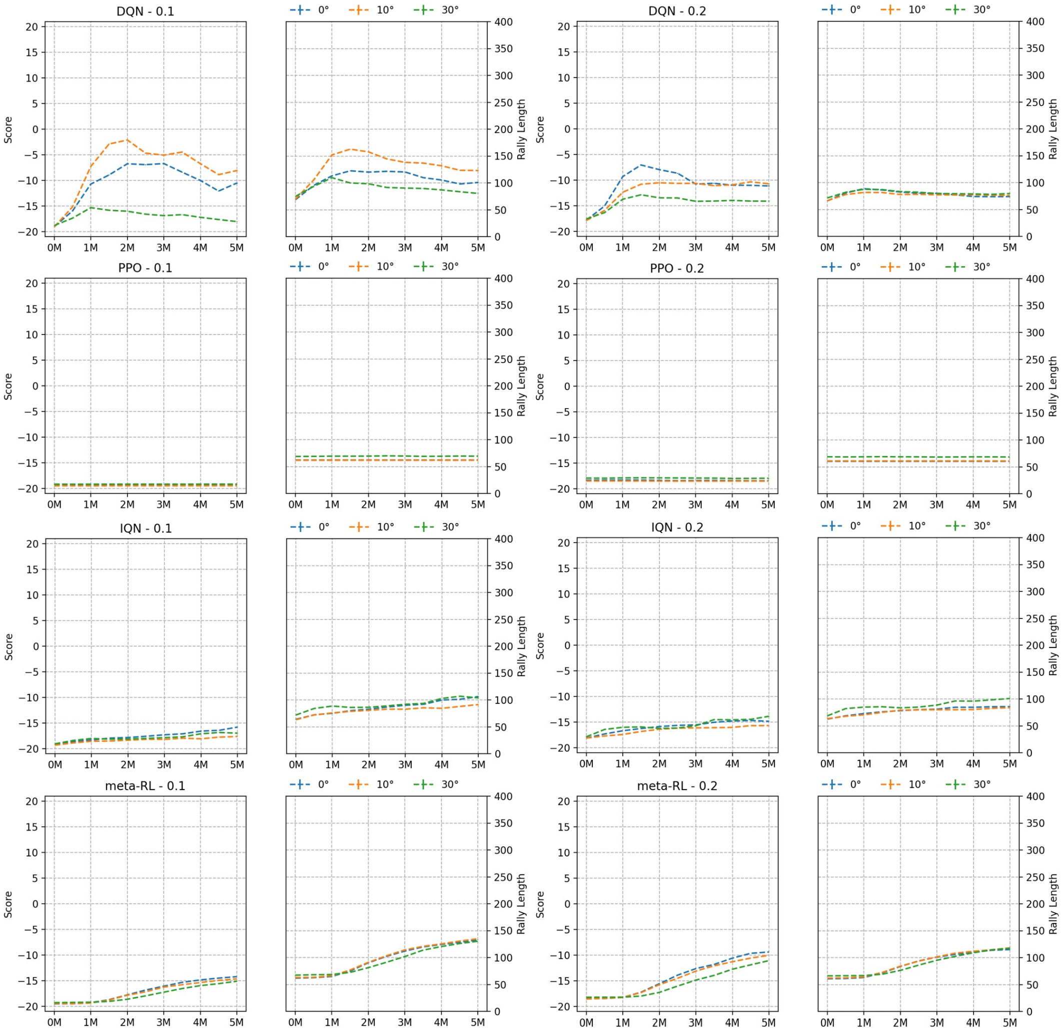

**Figure S1:** Deep RL Model Learning Summary
Blue (0°), Orange (10°), and Green(30°) for the LOW, MID, and HIGH uncertainty

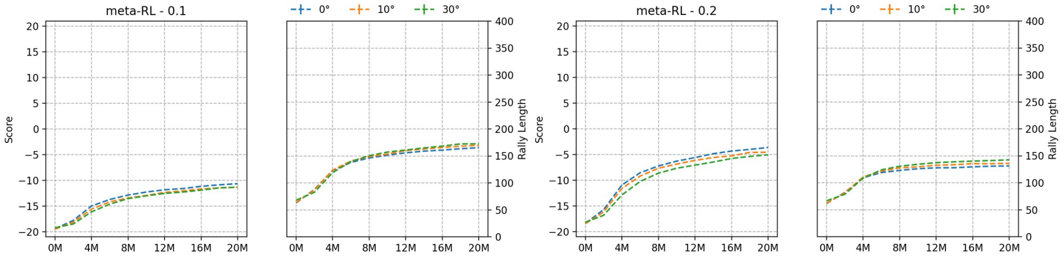

**Figure S2:** meta-RL Long-Term Learning
Blue (0°), Orange (10°), and Green(30°) for the LOW, MID, and HIGH uncertainty

Unlike the consistent gain shown in the 5M results (see figure S1), the learning of the meta-RL agent starts to decelerate after 5M interactions. Then, it converges to an asymptotic value, as shown in the figure S2. We did not present additional results (≥40M), as the change is miniscule, although the performance constantly improves.

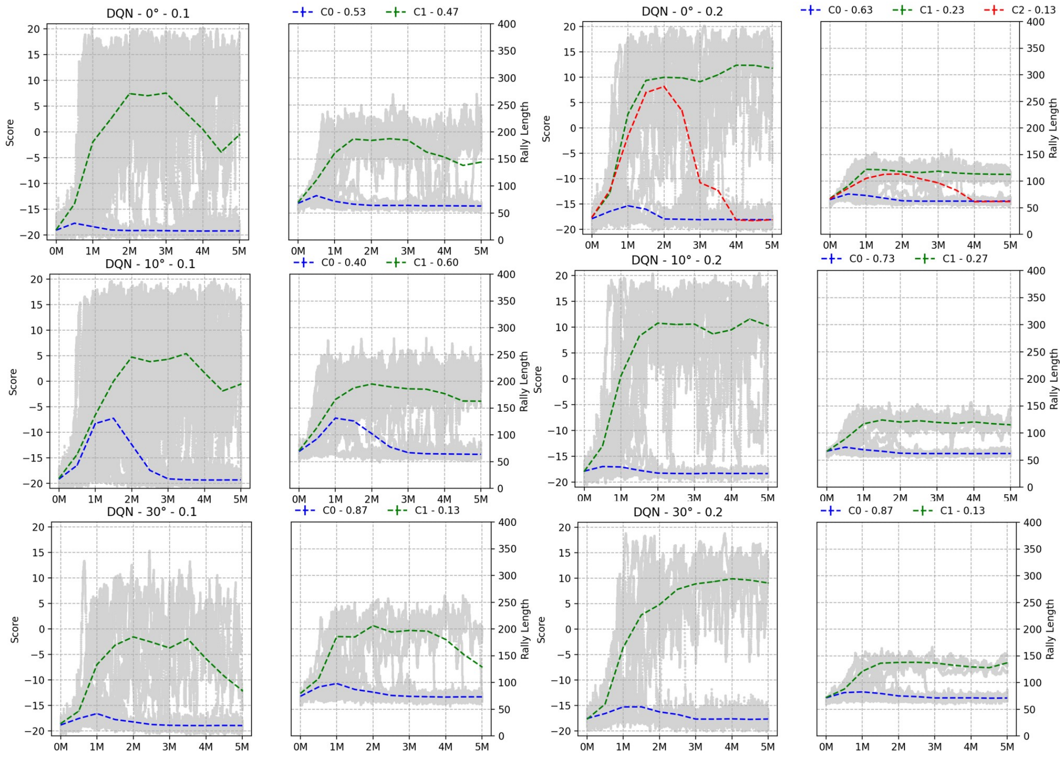

**Figure S3:** DQN Performance Bifurcation
Colored line for the average and s.e.m of the each group
The ratio of each group is in the right above for each task condition

The DQN agent learning branches out into 2-3 courses, depending on the task uncertainty and difficulty level (see figure S3). Initially, the DQN agent suffers from destructive forgetting under the HARD condition, even if the case agent learns successfully. Next, the ratio of successful learning increases when the task difficulty increases, except under the HIGH uncertainty condition. This indicates that the DQN agent learns dynamically from the interactions with the environment. In other words, if the quality of the supplied information increases, then it can learn more successful policy. However, this does not mean that it learns stably; all successful learning is suddenly impaired under the HARD condition, in contrast to the EASY condition. Third, under the HARD condition, the rally length increases and later decreases to a value similar to that in the EASY condition. This may suggest that the model capacity cannot handle the size of the environment state space. Finally, although the proportion is minute, a DQN agent in the EASY condition learns more stable policy. The other models do not exhibit such bifurcation, so we omitted their temporal clustering results.

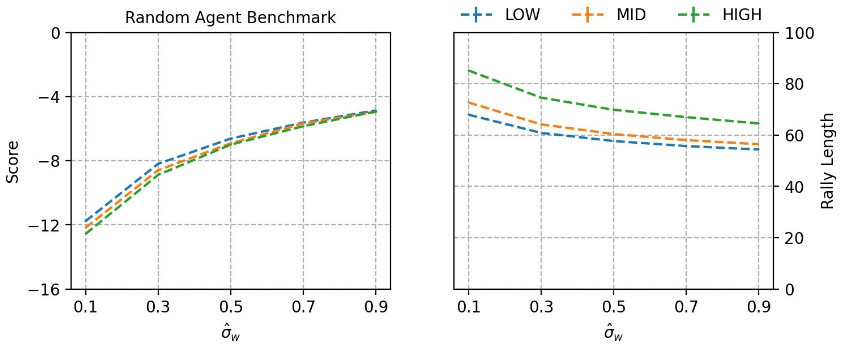

**Figure S4:** Random Agent Performance

The performance of the random agent keeps increasing when $\hat{\sigma}_W$ increases. However, the slope starts to decline (see figure S4). Additionally, the average rally length decreases because of the inflow of shorter rallies as the opponent will miss the ball more frequently when the ball is initially shot towards them. Furthermore, it converges to the number of frames required to horizontally traverse the table ($\simeq$60), which supports our view.

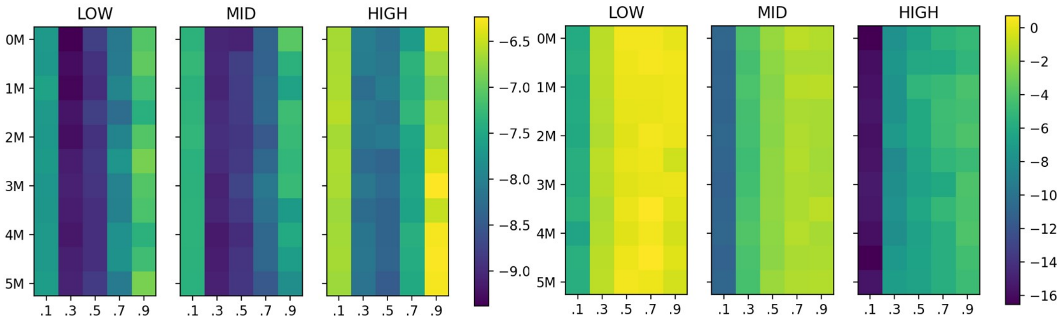

**Figure S5:** PPO Learning Deficiency
Left - Score Difference with Random Agent
Right - Rally Length Difference with Random Agent

Figure S5 shows the difference in the two measures between the PPO and the random agent. The consistent negativity of the score difference indicates that the PPO agent performs even worse than the random agent. However, when viewed horizontally, the gaps are continually narrowing, which means that the slope against $\hat{\sigma}_W$ of the PPO agent is larger. Moreover, when viewed vertically, for a difficulty level $\hat{\sigma}_W \geq 0.5$, performance improvement is observed. Therefore, we concluded that the PPO agent is simply unsatisfactory to tackle these tasks, but not incapable due to the errors.

##### 4.3.2 Ablation Study

**Table S4:** Ablation study results - score change [%] (5M episodes)

| Uncertainty | LOW | | MID | | HIGH | |
|---|---|---|---|---|---|---|
| Difficulty | EASY | HARD | EASY | HARD | EASY | HARD |
| TPP | -6.11 | -12.35 | -8.73 | -15.48 | -12.38 | -16.02 |
| SR | -84.07 | -90.08 | -83.82 | -89.07 | -81.56 | -87.73 |
| SSF | -50.71 | -74.96 | -46.87 | -72.03 | -46.60 | -67.37 |
| ELR | -4.56 | -5.83 | -3.52 | -4.73 | -5.39 | -9.08 |
| XOR | -61.33 | -75.84 | -61.15 | -75.63 | -59.82 | -74.45 |

**Table S5:** Ablation study results - rally length [%] (5M episodes)

| Uncertainty | LOW | | MID | | HIGH | |
|---|---|---|---|---|---|---|
| Difficulty | EASY | HARD | EASY | HARD | EASY | HARD |
| TPP | -50.81 | -37.61 | -50.80 | -31.13 | -48.14 | -33.01 |
| SR | -69.26 | -78.23 | -70.76 | -76.70 | -68.92 | -76.45 |
| SSF | -42.60 | -57.58 | -43.59 | -53.10 | -43.41 | -53.31 |
| ELR | -50.29 | -35.24 | -48.44 | -32.57 | -49.35 | -33.72 |
| XOR | -47.86 | -55.43 | -52.27 | -59.25 | -49.75 | -59.39 |

First, the entropy learning rate (ELR) is the only component for which the performance drops significantly when the uncertainty jumps from MID to HIGH. Under HIGH uncertainty, the most important step requires to the policy is establishing the primary division between events, where the decision should branch first. This division works as a primal context and helps the agent to solve such uncertain task. When considering the role of ELR, which dynamically allocates the learning rate to solve uncertainty, we can assume that it may work as a context manager. Second, the theta phase precession (TPP) consistently contributes to the learning, especially under the HARD condition. It serves the role that we will mention after.

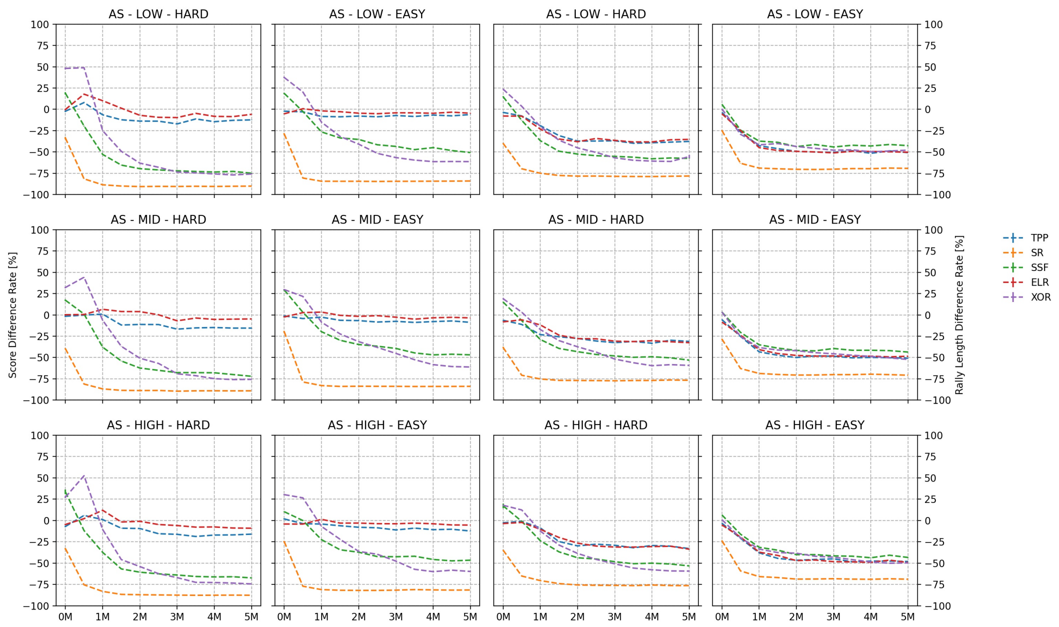

**Figure S6:** Ablation Study Summary
Difference rate $\gamma_{\Delta \mathrm{X}}$ is the difference $(\mathrm{X}_D - \mathrm{X}_N)$ divided by $\mathrm{X}_N$,
where $\mathrm{X}_D$ and $\mathrm{X}_N$ is the measure X of the deleted and the normal agent, respectively.

The last intriguing point is the role of the synaptic stability function (SSF). Unlike other deleted agents, the agent deleted SSF shows the decaying of the performance. The magnitude of the effect snowballs, and it loses all previously learned contents. However, the rally length remains constant, or even increases in some cases. From the tables S4-S5, the figure S6, and replay videos (data not shown), the agent fails to recall the policy at the moment immediately before missing a ball and proceeds to forget retrogressively along the time axis.

### 5 Analysis 2 - Biological Implication

#### 5.1 Additional Performance Measures for In-depth Analyses

Suppose that the Pong ball keeps moving toward the agent consistently, $\overrightarrow{v_x} > 0$. Until the ball finally reaches $x_B$ on the pad line, we store (1) the firing activity of all PFC cells $\mathbf{A}_P \in \mathbb{R}^{L_T \times N_P}$ and (2) the most probable reward position predicted by the agent every time $\mathbf{x}_A^* \in \mathbb{R}^{L_T \times 1}$, where $L_T$ is the length of the records. The oldest record is located in the first row, and the newest record is located in the last row. From these raw data, we can obtain the time coordinate $\mathbf{t}_R$ and two measures $\mathbf{E}$ and $\mathbf{R}_C$, denoting the error and the accuracy, respectively.

##### 5.1.1 Relative Time

The definition of relative time $\mathbf{t}_R \in \mathbb{R}^{L_T \times 1}$ is the arithmetic sequence starting from 0 (oldest) to 1 (newest) allocated to each row of $\mathbf{A}_P$ and $\mathbf{x}_A^*$.

$$\mathbf{t}_R[k] = \frac{k-1}{L_T - 1} \qquad \text{for } \; k = 1, 2, \ldots, L_T$$

##### 5.1.2 Timing Error

To hit the approaching ball, the agent must move its panel. At each instant, we can calculate the time required for the agent to move and defend successfully from the current position. In other words, this time is the estimated number of frames the agent moves and confines the ball inside the range covered by its panel:

$$\mathbf{t}_A = \left\lceil \frac{\min\left(\left|\mathbf{x}_A^* - x_B\right| - W_A, 0\right)}{V_A} \right\rceil$$

where $W_A$ is the half width, and $V_A$ is the speed of the Pong panel. For each row of the record, remaining time exists until the end $\mathbf{t}_O = [L_T, \ldots, 2, 1]^T$. The error $\mathbf{E} \in \mathbb{R}^{L_T \times 1}$ for each frame is

$$\mathbf{E} = \arctan\left[\frac{\mathbf{t}_A}{\mathbf{t}_O}\right]$$

##### 5.1.3 Cell Population Accuracy

We define the accuracy $\mathbf{R}_C$ as the proportion of the "correct" PFC cell. Here, the "correctness" means that its firing, a tool to vote for upside when fires, matches the relative direction between the actual position at which the ball arrives and its decision center. The accuracy $\mathbf{R}_C \in \mathbb{R}^{L_T \times 1}$ is

$$\mathbf{F} = u\left(x_B - \mathbf{x}_P\right) \qquad \mathbf{R}_C = \frac{\overline{\mathbf{A}_P \oplus \mathbf{F}}}{N_P} = \frac{\left[\mathbf{A}_P \cdot \mathbf{F} + \left(J_{L_T, N_P} - \mathbf{A}_P\right) \cdot (\mathbb{1} - \mathbf{F})\right]}{N_P}.$$

where $J$ is a matrix of ones. The operation above is essentially the equivalence operation.

##### 5.2 Experimental Procedure

First, to test the contributions under harsher conditions, we changed the number of PFC cells $N_P$ from 100 to 20 and the half width of Pong pad from 14 to 7. In this way, the agent needs to manage the same tasks with much more confined resources, forcing the agent to increase the efficacy of its policy. The epochs are identical to 50 games ($\simeq$150K interactions) for all simulations. After each game, all the agents conduct 60 test games, composing one test session for each training game.

Then, we discretized the relative time $\mathbf{t}_R$ into 40 bins and computed the average of the difference of measures between two groups at the last relative time bin of the individual test session. We considered the last relative time bin because the discrepancy as well as the contribution of each function are highlighted most at the moment just before the arrival of the Pong ball. Furthermore, we classified tests into two cases depending on whether the agent hits the ball successfully for the comparison. The measures used are the error $E$ and the accuracy $R_C$.

###### 5.2.1 Theta Phase Precession

After preprocessing the data, we analyzed how $\Delta E$ evolves while the agent learns the task. To do this, we smoothed the raw data using simple exponential smoothing after the optimization. Then, we clustered 50 data points into 5 groups and calculated the averages and the standard errors of the means for each group (containing 10 data points). We repeated this procedure for all six task conditions.

###### 5.2.2 Successor Representation - Update Distance

After preprocessing the data, we analyzed how $\Delta E$ at last test session varies while $l_U$ changes continuously. To do this, we smoothed the raw data using simple exponential smoothing after the optimization. Then, we clustered 50 data points into 5 groups and calculated the average and the standard error of the means for the last group alone. We repeated this for all six task conditions.

Furthermore, to investigate the trend accurately, we optimized the fitting hyperparameter, the upper boundary $l_U^*$, which the performance becomes destructed if $l_U \geq l_U^*$. We performed the grid search and obtained the argument of the maxima in terms of the total sum of log-likelihood of the fitting result. This idea will be discussed in detail in a subsequent section.

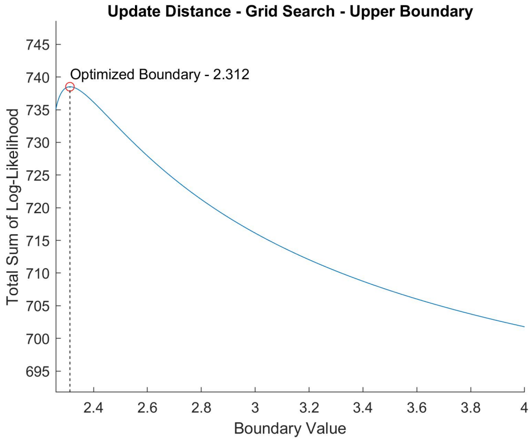

**Figure S7:** Hyperparameter Search
The optimization result of the upper boundary $l_U^*$ by grid search.

###### 5.2.3 Successor Representation - Current Distance

After preprocessing the data, we analyzed how the offset of $\Delta R_C$ varies while $l_C$ changes continuously. To do this, we smoothed the raw data using simple exponential smoothing after the optimization. Then, we calculated the average and the standard error of the means of all 50 data points. We repeated this procedure for all six task conditions.

### 6 Role of Predictive Spacetime Encoding in Learning

Assume that our learning model is a theoretical system following the information theory. Then, the spacetime SR module serves the role of an encoder, with each component contributing distinctively. When we consider the biological place cell as an abstract symbol or character, the engagements of each composing mechanism become highlighted.

#### 6.1 Theta Phase Precession

The theta phase precession expands the dimension of representation, now capable of weighting differently when transferring the signal to the decoder. It gains two advantages if the agent achieves abstract policy that appreciates the future symbols most; the decoder can make a context-dependent decision as different future events direct to different contexts. Further, the agent can allocate more importance to the future, the role of support, or guidance for the past, while learning reciprocally.

#### 6.2 Successor Representation

Next, the successor representation (SR) modulates the symbol space. In our model, SR first pre-allocates or tunes the receptive field of a single symbol to focus on the relevant symbols. Next, it combines the receptive fields of the current symbols and signifies the overlapping ones. We divide this modulation into two compartments, one for each hyperparameter.

##### 6.2.1 Update Distance

The update distance $l_U$ regulates the number of other symbols that a single symbol should represent. If ensuring the right size of the current distance $l_C$, a small $l_U$ will not matter as the SR matrix now encodes the context using a combination of the current symbols rather than overlapping. In contrast, a large $l_U$ exacerbates the problem. Because each receptive field is excessively blurred, the overlapping marks entire adjacent cells, including the irrelevant to the context, and hinders the decision.

If there is no uncertainty, only one right decision and context exists, while the others are completely incorrect. With such a sharp decision boundary over the contexts, this obscurity by overlarge $l_U$ leads to a critical deficiency. On the other hand, there are less distinct right decisions and wrong decisions if the agent resides in a task with high uncertainty. Because there is innate blurring caused by the environment, any additional dimness only slightly degrades the performance. Therefore, we can infer that this issue becomes critical when the task uncertainty declines due to the sharpness. To prove this, we compared the error $E$ between the agent with shifted $l_U$ and the agent with normal, fixed $l_U$. When we increase this shifted update distance, $\Delta E$ should start to drop at a certain point and become nosedived over a specific boundary. Moreover, such lessening should increase when the task uncertainty decreases.

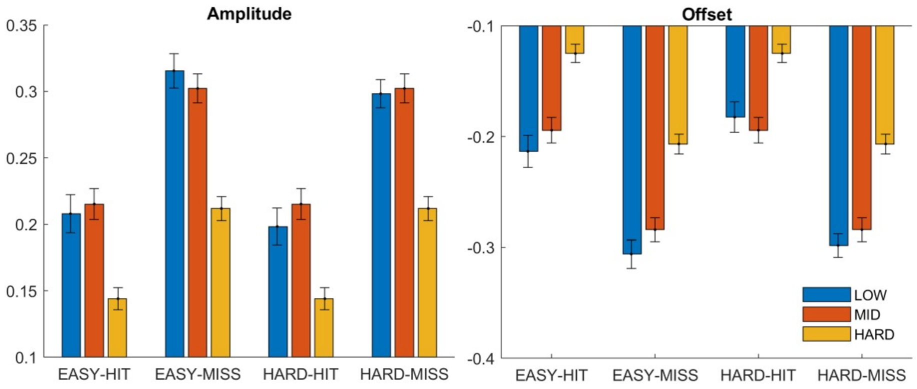

**Figure S8:** Update Distance and Error Difference
Fitting with $\Delta E = A \cdot \sqrt{l_U^* - l_U} + C_{\text{OFF}}$, where $l_U^*$ is the fitted upper boundary of $l_U$ (see figure S7). $l_U^*$ corresponds to the value of the red line in the figure S9 and S10

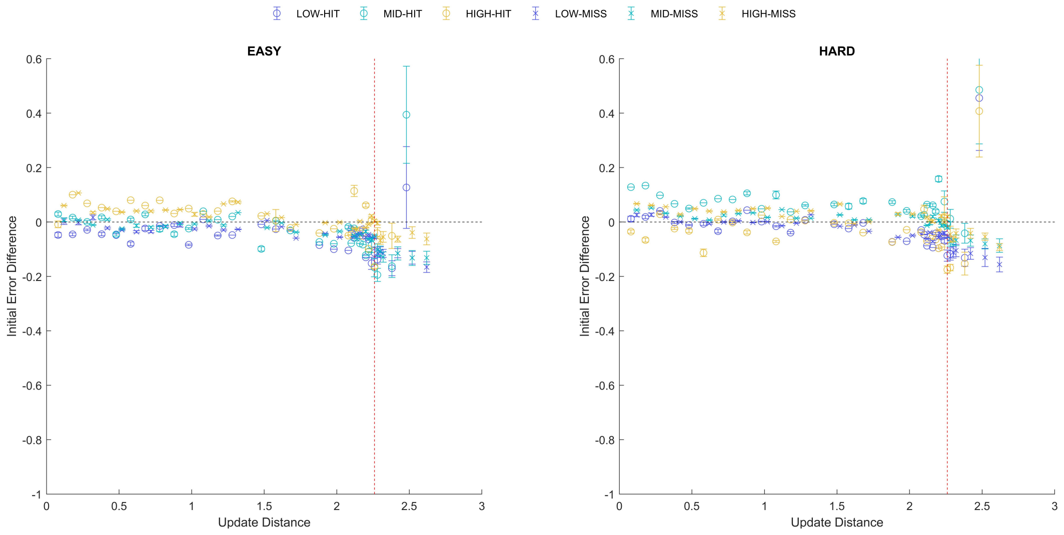

**Figure S9:** Initial Error Difference
Initially, the update distance does not affect significantly.
The data points are the densest along the $\Delta E = 0$

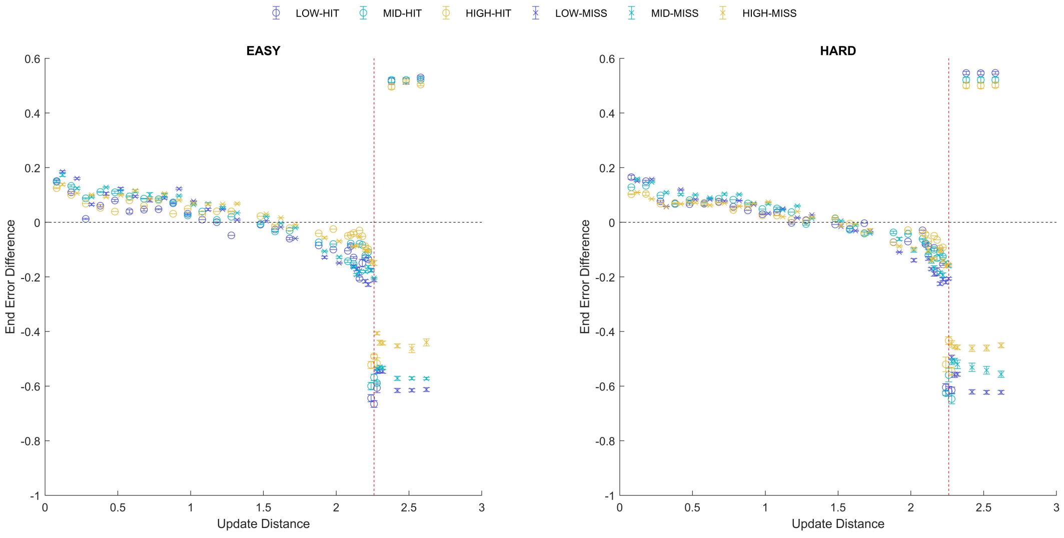

**Figure S10:** Terminal Error Difference
Although the starting points are similar, the results of learning differ significantly.
Beyond the red line $l_U^*$, the agent mostly makes wrong decisions (≥90%) with huge errors.

As shown in figure S9 and S10, $\Delta E$ values for small $l_U$ are almost identical for all uncertainty levels. However, $\Delta E$ curves differently as $l_U$ increases, and it drops permanently when $l_U$ exceeds the red line (see figure S10), although this decrease does not appear in the initial period (see figure S9). Moreover, the instantaneous rate of decrease of $\Delta E$ (see figure S8 Amplitude), and also the terminal value just before the asymptotic value, the red line, (see figure S8 Offset) becomes more negative when the uncertainty decreases, as we predicted.

##### 6.2.2 Current Distance

The current distance $l_C$ modulates the number of symbols that fire simultaneously. In other words, it is theoretically equivalent to the source coding theorem in information theory. Generally, the accuracy of the learning system increases and slowly converges to the certain value. Here, there are two quantities we need to consider: the accuracy after complete learning, and the number of total interactions required to achieve this accuracy. Suppose that there is an optimized current distance $l_C^*$ that can support a sufficient number of symbols to represent the entire environment.

First, we start with the accuracy after saturation. If $l_c \leq l_C^*$, the accuracy $R_C$ of the decoder degrades because there are events that the encoder cannot represent. However, in the opposite case, $R_C$ declines due to abundance of unnecessary symbols, leading to the noisy channel.

Next, consider the number of interactions until the saturation. The larger the size of the symbol space becomes, the longer the time required to learn the policy. This is because the decoder unit needs to differentiate more symbols and construct more complex decision boundaries.

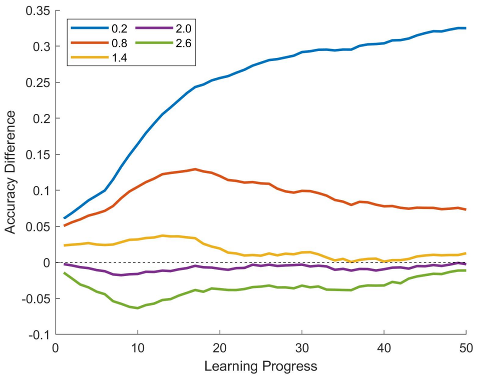

**Figure S11:** Accuracy Difference during Learning
Learning Progress VS Accuracy Difference for Different $l_C$

From the above statements, we can infer two properties. If we constrain the period of learning, the average $\Delta R_C$ should be positive for small $l_C$ and continue to drop when $l_C$ increases due to increased saturation time (see figure S11). Furthermore, due to the source coding theorem, the source (environment) with higher uncertainty requires more symbol to represent entire events. Subsequently, it means that although two agents have same $l_C$ (same number of symbols), the agent under lower uncertainty can achieve higher accuracy than the agent under higher uncertainty. Therefore, the increasing uncertainty should decrease the saturated accuracy for same $l_C$. (see figure S12 Amplitude).

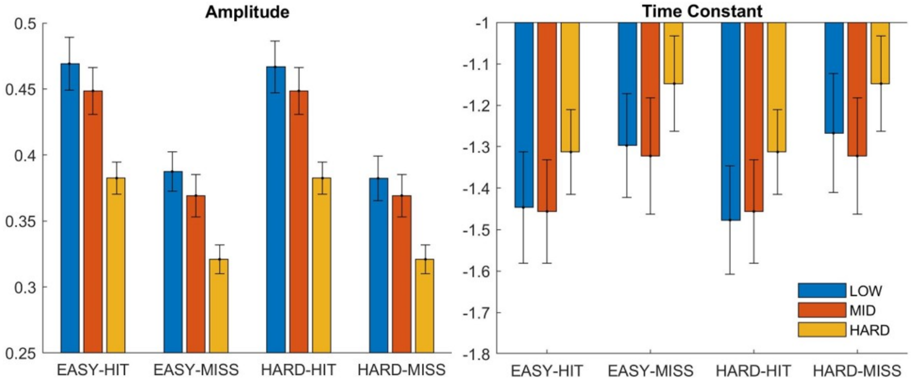

**Figure S12:** Current Distance and Average Accuracy Difference
Fitting with $\Delta R_C = A \cdot \exp\left[\tau_C \cdot l_C\right]$

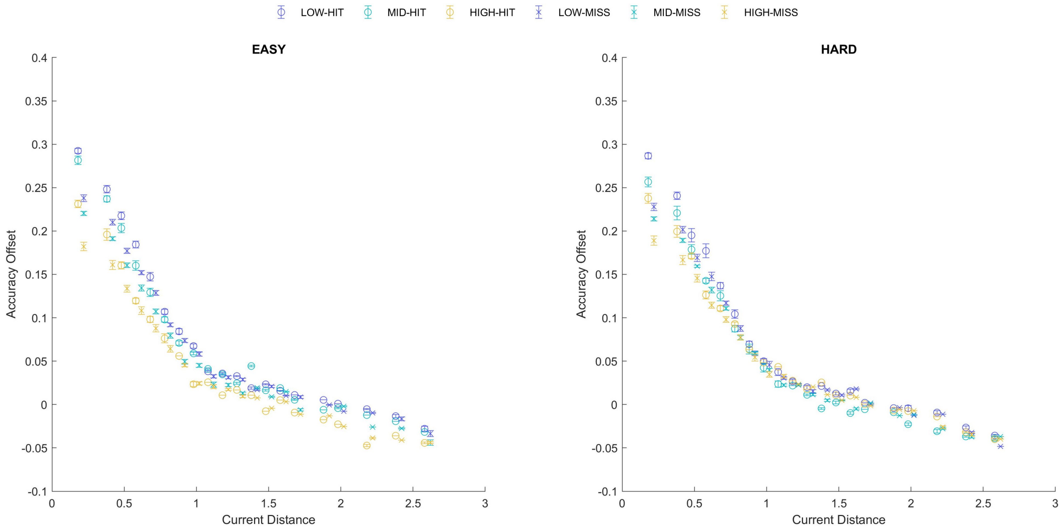

**Figure S13:** Average Accuracy Difference

As shown in the fitting result (see figure S12) and raw data (see figure S13), these figures support our statements regarding the uncertainty and difficulty level of the task. In addition, we observed that the instantaneous rate of decrease of $\overline{\Delta R_C}$ decreases when the uncertainty level increases (see figure S12 Time Constant). In fact, these effects are significant when the agent learns from a few interactions. When we increase the number of interactions to ($\geq$1M), such effects disappear as the accuracy saturated completely in the moderate range of $l_C$ (data not shown). Subsequently, the accuracy offset becomes in the shape of the parabola with the negative focus point, centered at the $l_C^*$.

[7] Volodymyr Mnih, Koray Kavukcuoglu, David Silver, Alex Graves, Ioannis Antonoglou, Daan Wierstra, and Martin Riedmiller. Playing atari with deep reinforcement learning. *arXiv preprint arXiv:1312.5602*, 2013.

[8] Ashley Hill, Antonin Raffin, Maximilian Ernestus, Adam Gleave, Anssi Kanervisto, Rene Traore, Prafulla Dhariwal, Christopher Hesse, Oleg Klimov, Alex Nichol, Matthias Plappert, Alec Radford, John Schulman, Szymon Sidor, and Yuhuai Wu. Stable baselines. `https://github.com/hill-a/stable-baselines`, 2018.

[9] Hado Van Hasselt, Arthur Guez, and David Silver. Deep reinforcement learning with double q-learning. In *Thirtieth AAAI conference on artificial intelligence*, 2016.

[10] Ziyu Wang, Tom Schaul, Matteo Hessel, Hado Van Hasselt, Marc Lanctot, and Nando De Freitas. Dueling network architectures for deep reinforcement learning. *arXiv preprint arXiv:1511.06581*, 2015.

[11] John Schulman, Filip Wolski, Prafulla Dhariwal, Alec Radford, and Oleg Klimov. Proximal policy optimization algorithms. *arXiv preprint arXiv:1707.06347*, 2017.

[12] Will Dabney, Georg Ostrovski, David Silver, and Rémi Munos. Implicit quantile networks for distributional reinforcement learning. *arXiv preprint arXiv:1806.06923*, 2018.

[13] Jane X Wang, Zeb Kurth-Nelson, Dhruva Tirumala, Hubert Soyer, Joel Z Leibo, Remi Munos, Charles Blundell, Dharshan Kumaran, and Matt Botvinick. Learning to reinforcement learn. *arXiv preprint arXiv:1611.05763*, 2016.
